# Factors affecting virus prevalence in honey bees in the Pacific-Northwest, USA

**DOI:** 10.1101/2021.11.19.469300

**Authors:** Vera W Pfeiffer, David W Crowder

## Abstract

Global efforts to assess honey bee health show viruses are major stressors that undermine colony performance. Identifying factors that affect virus incidence, such as management practices and landscape context, could aid in slowing virus transmission. Here we surveyed viruses in honey bees from 86 sites in the Pacific Northwest, USA, and tested effects of regional bee density, movement associated with commercial pollination, julian date, and hive management on virus prevalence. We also explored patterns of virus co-occurrence and spatial autocorrelation to identify whether local transmission was a primary driver of pathogen distribution. Our surveys found widespread prevalence of Deformed wing virus (DWV), Sacbrood virus (SBV), and Black queen cell virus (BQCV). BQCV and SBV were most prolific in commercial apiaries, while Chronic bee paralysis virus (CPBV) was more common in hobbyist apiaries than commercial apiaries. DWV was most common in urban landscapes and was best predicted by mite prevalence and julian date, while the incidence of both SBV and BQCV were best predicted by regional apiary density. We did not find evidence of additional spatial autocorrelation for any viruses, although high co-occurrence suggests parallel transmission patterns. Our results support the importance of mite management in slowing virus spread and suggest that greater bee density increases transmission. Our study provides support that viruses are widespread in honey bees and connects known mechanisms of virus transmission to the distribution of pathogens observed across the Pacific Northwest.

**Highlights:** Three viruses were widespread in honey bee populations across the Pacific Northwest, USA

Black queen cell and Sacbrood viruses were most common in high density hives

Deformed wing virus was most common in hives that had high mite loads

The presence of many viruses in bees suggests parallel or synergistic transmission

## Introduction

The health of honey bees is a global economic and ecological concern, as worldwide movement of biotic materials promotes the spread of pathogens and pests which adversely affect bee health. Indeed, at least 24 viruses are known to cause disease in honey bees (Brutscher et al., 2016; Chen and Siede, 2007). Movement of honey bee apiaries to meet pollination demands of fruit and nut crops is also cited as a major concern for virus spread, as virus transmission occurs through close contact among nestmates, and when infected bees drift into other colonies (Dynes et al., 2019). Such conditions that promote virus spread may be most prevalent in areas where honey bee apiaries are stocked at high densities to meet pollination needs. However, while multiple factors can increase bee exposure and susceptibility to viruses, the most consequential factors determining virus transmission and susceptibility across variable landscapes are often unclear.

While apiculture and domesticated honey bee populations continue to grow worldwide, honey bee stocks are increasing at a rate slower than the demand for agricultural pollination (Aizen and Harder, 2009). Several studies show higher virus incidence in landscapes with crops that rely on commercial pollination compared to those without commercial pollination (Alger et al., 2019; Olgun et al., 2020). While much of the focus on honey bee health has assessed rural ecosystems where commercial apiaries are managed for agricultural pollination, urban ecosystems have also seen rapid growth in the number of hobbyist beekeepers that maintain hives for personal gardens. Improved knowledge of virus prevalence in both rural and urban ecosystems can support activities to prevent virus introduction into non-infected regions or apiaries, or spread between colonies within apiaries. Virus mitigation can also be attempted by controlling other pathogens that may act in synergy, although it is often unclear if different viruses are transmitted concurrently or independently from one another (Aubert et al. 2011).

Recent surveys suggest that not only are viruses more prevalent than previously known, but co-occurrence of viruses in single colonies is common, and that honey bees are more susceptible to secondary infection once infected (D’Alvise et al., 2019). Viruses may be pathogenic alone, but pathogenicity may be induced by other factors including hunger, cold, toxicants, or other pathogens (Doublet et al., 2015, Di Prisco et al., 2013, Dolezal et al., 2019). Relative occurrence rates of pathogens often differ by region and pathogen type, and weak and declining colonies may become susceptible to an array of pathogens. Moreover, the synergistic effects of multiple pathogens deplete workers and lead to more frequent colony demise (Cornman et al., 2012; Burnham et al., 2019). However, few studies have conducted virus sampling across broad enough regions, and at enough sites, to determine the spatial autocorrelation among pathogens that may provide evidence of parallel transmission patterns.

In this study, we aimed to investigate how known factors related to virus transmission and virulence explained the distribution of honey bee viruses at a broad landscape scale, and what geographical patterns may result from the manifestation of these relationships. We predicted that co-infection of multiple viruses is more common than expected based on virus prevalence due to synergistic effects between viruses (D’Alvise et al., 2019). We also hypothesized that increasing regional bee density, greater apiary movement associated with commercial pollination, and lack of mite treatments may drive increased prevalence of bee viruses due to increased transmission or greater bee susceptibility. Consequently, we expect to notice more virus prevalence in regions with high density of apiaries and high use of commercial pollination. Our study was conducted on over 80 sites across a broad region encapsulating both urban, agricultural, and rural ecosystems, giving us sufficient power to tease apart these relationships.

## Materials and Methods

### Bee Sampling

We collected 30 honey bees from each of 86 sites (n = 2,580 bees) across Washington state and adjacent parts of Oregon and Idaho (Fig. 1). These sites reflected various landscape types including urban, agricultural, mixed-use residential, forested, and steppe. Sampling occurred between July 10th and August 28th, 2020. Sixty-eight sites had active apiaries; the other 18 sites had honey bees foraging but no visible apiary. For the sites with apiaries, foraging honey bees entering and leaving apiaries were netted until 30 were collected. At sites without apiaries (e.g. urban community gardens), 30 honey bees were sampled by hand net. Apiary management surveys (Table S1) were collected from 54 participating beekeepers, including 5 sites with commercial apiaries and 49 hobbyist beekeepers with less than 20 hives. We were not able to obtain completed surveys from the other 14 sites with apiaries. Netted bees were deposited in 5ml centrifuge tubes and euthanized in dry ice in the field, then stored at -20°C until cataloged, and then stored at -80°C until RNA extraction. Nets were sanitized between sites.

**Figure 1.**
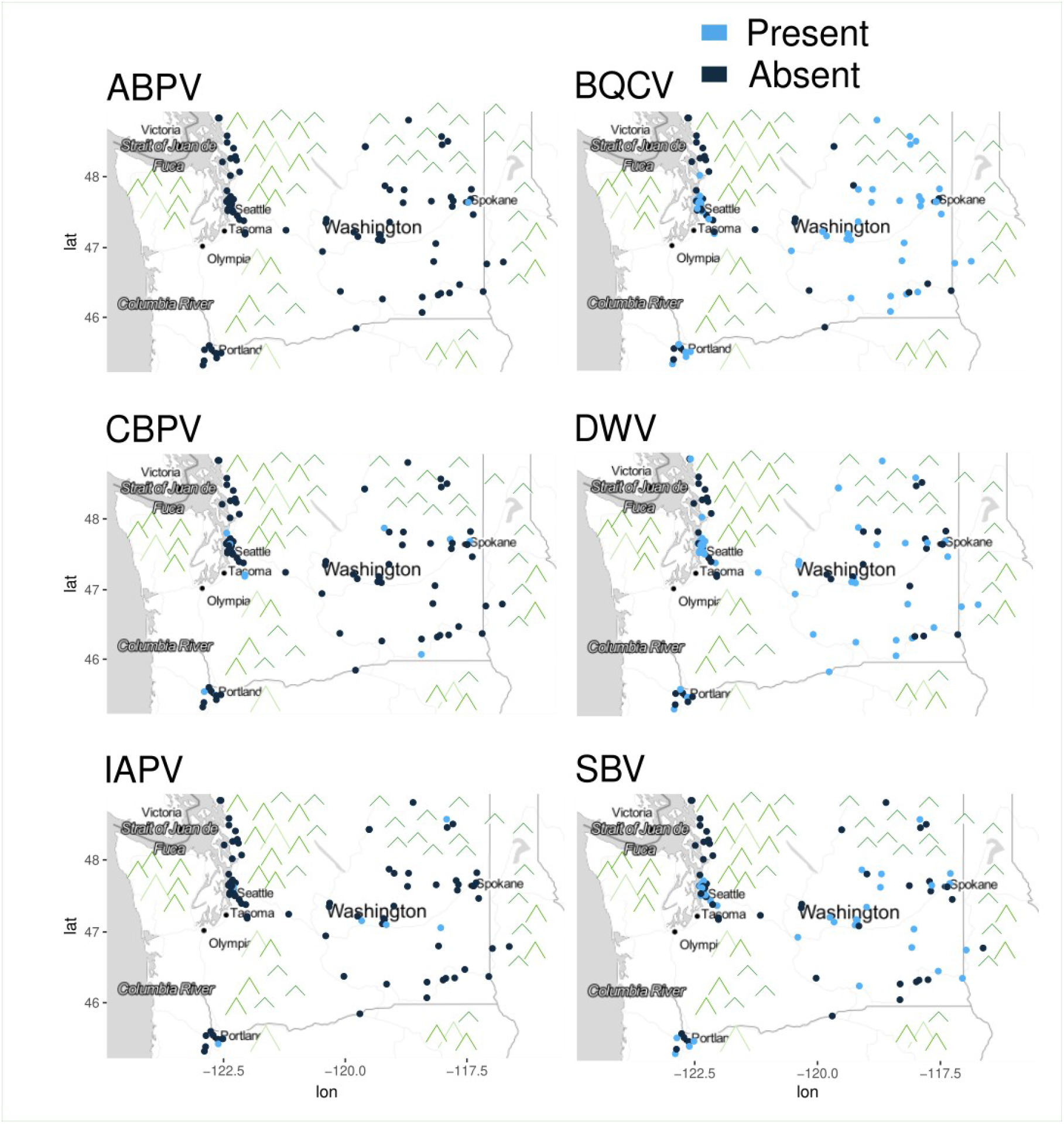
Maps of Acute bee paralysis virus (ABPV), Black queen cell virus (BQCV), Chronic bee paralysis virus (CBPV), Deformed wing virus (DWV), Israeli acute paralysis virus (IAPV), and Sacbrood virus (SBV) incidence at 86 sampling locations spanning between three cities in the northwestern USA – Seattle, WA, Spokane, Washington, and Portland, Oregon.

### Viruses Assessed

Honey bee management for bee products and agricultural pollination is a global occupation, and most common bee viruses are observed around the world (Goulson and Hughes, 2015). While several viruses manifest with unique observable symptoms, most are also found as asymptomatic infections (Grozinger and Flenniken, 2019). However, increased efficiency of molecular diagnostic methods has improved the capacity for rapid and widespread virus detection. In this study we used molecular methods to test for several viruses described here.

Sacbrood virus (SBV) was the first honey bee virus identified as the pathogen responsible for liquifying larvae, and has recently been considered the most widely distributed honey bee virus (Chen and Siede, 2007; White et al., 1913). While larvae are most susceptible to SBV, infected adults may have a decreased life span (Bailey, 1969). SBV is spread within the colony when nurse bees become infected while removing infected larvae and then they transmit the virus while feeding larvae and exchanging food with other bees (Chen and Siede, 2007). SBV infection thus arises seasonally in the summer with the proliferation of susceptible brood.

Deformed wing virus (DWV) was first isolated in Japan, and subsequently has been found around the world. Deformed wing virus can be asymptomatic but also can cause shrunken and crumpled wings, reduced activity, decreased body size, and increased mortality. Adverse impacts have been recorded in bumble bee species as well as *Apis mellifera*. DWV is known to be transmitted by trophallaxis and shared food resources, as well as *Varroa destructor* mites, whose abundance is strongly correlated with winter losses (Chen and Siede, 2007; Grozinger and Flenniken, 2019; Yang and Cox-Foster, 2007).

Black queen cell virus (BQCV) was first isolated from dead queen larvae and prepupae sealed into dark brown cells (Bailey and Woods, 1977), and is frequently the most common honey bee virus reported from North America and Europe. Larvae may exhibit pale yellow coloration and saclike skin similar to SBV infected larvae. Infected workers do not exhibit symptoms, and the virus does not tend to multiply in bees after ingestion. BQCV infection is associated with *Nosema apis* infection, where BQCV multiplies rapidly in the bee’s body when infected with the *Nosema apis*, fungal pathogen (Bailey et al., 1981; Bailey and Perry, 1982). Infection may also be associated with *Varroa destructor* (Tentcheva et al., 2006, 2004).

Three less common viruses assessed were chronic bee paralysis virus (CBPV), acute bee paralysis virus (ABPV), and Israeli acute paralysis virus (IAPV). CBPV was identified as a cause of adult bee paralysis in 1963 (Bailey et al., 1963), and field surveys of mites show they do not transmit the virus. ABPV was discovered during lab infectivity tests of CBPV, and replicates faster than CBPV (Chen and Siede, 2007). ABPV was originally considered an economically irrelevant virus in honey bees, however, both brood and adult bee mortality were later observed in colonies infested with *Varroa destructor* (Grozinger and Flenniken, 2019). ABPV may also be triggered by other causal factors (Chen and Siede, 2007). IAPV is a more recently described virus, that has been associated with shivering wings, progressing to paralysis, and death of workers outside the hive, as well as colony collapse disorder symptoms, and may also be spread by Varroa destructor mites (Cox-Foster et al., 2007, Di Prisco et al., 2011; Maori et al., 2007).

### Bee virus assessment

To assess viruses, the 30 honey bees from each site were divided into 3 groups of 10. With this scheme we had 258 total samples (86 sites × 3 groups of 10 honey bees per site = 258), although one sample was destroyed during processing, resulting in 257 samples analyzed in total. Honey bee thoraxes were isolated from each bee; heads and abdomens that contain inhibitory enzymes and compound eyes were separated and removed (Boncristiani et al., 2011). RNA was extracted from bee thoraxes from each site and pooled for each group of 10 bees. The ten thoraxes that made up each sample were placed in a nuclease-free centrifuge tube (2ml), then glass beads and Trizol (1ml per tube) were added before homogenization in the BeadRupter for two 30 second intervals at 4m/s and 6m/s. Following homogenization, 200ul of chloroform were added and tubes were vigorously vortexed for 15 sec, then allowed to sit on ice for 15 min. After settling, samples were centrifuged at 14,000 gravity (g) for 20 min. The aqueous phase was then transferred into a fresh tube, and isopropanol (0.5ml per ml of TRIzol) was added and mixed by inverting the tube. Samples were left on ice for 40 min, then centrifuged at 14,000 g for 10 min to precipitate and separate the RNA in a small pellet. RNA pellets were washed with 1 ml 75% ethanol twice, and centrifuged at 7,500 g for 5 min. The ethanol was poured off and pellets were allowed to air dry before resuspending in 1 ml nuclease-free water and stored at -80 °C. The concentration of the extracted RNA was measured on a Nanodrop 2000c (Thermo Fisher Scientific, Waltham, MA).

Complementary DNA (cDNA) was synthesized through reverse transcriptase PCR. 1ug of RNA diluted in 16 μl of water and 4ul cDNA iScript master mix (Promega, Madison WI) were combined in a 20 ul reaction. The cDNA was synthesized in a thermocycler program: one cycle at 94 °C for 5 min followed by 56 °C for 30 s, and 72 °C for 45 s. cDNA products were stored at -20 °C. We then used multiplex RT-PCR to detect the six bee viruses in a 25 μl reaction with 0.5 ul of each of the 10 mM oligonucleotide primers, 12.5 Taq mastermix (supplied with enzyme) and 1.5 μl of cDNA. Multiplex RT-PCR is an efficient and sensitive technique for simultaneous detection of different viruses in a sample; while the method does not characterize individual sequences it allows for detection of variants of individual viruses as long as there is no mutation in the primer annealing site. Multiplex-PCR was conducted using the following parameters: one cycle at 94 °C for 5 min followed by 35 cycles at 94 °C for 30 s, 56 °C for 30 s, and 72 °C for 45 s and a final extension cycle at 72 °C for 10 min. PCR products were analyzed by electrophoresis on a 1.5% agarose gel (100 V for 60 min). After completing the analyses, we spiked eight PCR reactions with cDNA from four known positive viruses and observed positive amplification in each reaction, implying the multi-plex was capable of detecting individual viruses effectively.

### Measuring factors that may affect virus spread

Participation in the study was requested via several associations: the Washington Beekeepers, the Portland Beekeepers, the Puget Sound Beekeepers, and the Mid-Columbia Beekeepers, as well as Backyard Beekeepers of Spokane, WA. Bees were sampled from all respondents who maintained contact following our initial request. Volunteer beekeeper participants who provided hives for testing also provided data on factors used in the statistical analysis. First, regional bee density was coded as a ranked value of 1 to 4, 1 indicated 0 or 1 known apiary in the surrounding 10km, 2 indicated 2-5 known apiaries in the surrounding 10km, 3 indicated 5-10 known apiaries in the surrounding 10km, and 4 indicated > 10 known apiaries or any large commercial pollination use within the surrounding 10 km. We also collected data on whether hives were moved during the year (yes or no), whether any disease treatments were used (yes or no), and whether mites were present in hives (yes or no). We recorded the julian date (ordinal date) of sampling to represent the hypothesis that viruses prevalence increases during the summer with increased population size and activity.

#### Statistical analysis

To test our hypotheses that bee density, bee movement associated with commercial pollination, mite presence, julian date, and mite treatments predicted virus incidence, we used generalized linear mixed models fit by maximum likelihood (Adaptive Gauss-Hermite Quadrature to approximate the log-likelihood) using the 54 sites from which we obtained management surveys. Fixed effects represented explanatory variables, and a random effect was included to represent the apiary site. We assessed whether common bee viruses are more prevalent in commercial apiaries, and certain apiary rich landscapes and ecotypes using contingency tables depicted with mosaic plots. We used Chi square tests and Fisher’s exact tests to identify significant differences in virus prevalence across categories. Subsequently, we investigated the role of additional spatial autocorrelation in our virus dataset using spatial regression. We averaged the three quantified band brightness virus estimates from each site across the full dataset of 86 sites, created a list of neighbors using the Queen criteria, generated the spatial weights matrix, and applied the Moran’s test on regression residuals in preparation to fit a spatially lagged regression model, which was finally not justified based on the lack of significance of the Moran’s test.

## Results

We collected thirty honey bees from each of the 86 sites that included 18 commercial apiaries, 50 hobbyist apiaries, and 18 other sites (Fig. 1). Of the surveyed apiarists, 76% of beekeepers reported mites. Each apiary with over 20 hives used chemical and cultural mite control. Fourteen percent (n = 7) of small apiary beekeepers had not used chemical treatment for mites by the time bees were sampled in July or August, and 12% (n = 6) opted for no disease treatments.

### Virus prevalence across the study extent

Of the 257 samples processed, 178 tested positive for at least one virus (69%) (Table 1). Three viruses were broadly distributed, BQCV observed in 97 positive tests from 52 of 86 sites (60%), DWV observed in 92 positive tests from 47 of 86 sites (55%), and SBV observed in 65 positive test results from 36 sites (42%). The sparsely observed viruses, ABPV, CBPV, and IAPV were only observed at 1, 12, and 6 sites, respectively.

**Table 1.** (A) Incidence and prevalence of viruses by samples (n = 257) and study sites (n = 86). The ‘sample incidence’ indicates the number of samples where viruses were observed out of the total 257 samples tested (86 sites × 3 samples per site, with one sample destroyed). This variable differs from ‘site incidence’, which indicates the number of sites (out of 86) that had a least one sample testing positive (with 3 pools of honey bees tested per site). (B) Pearson correlations between viruses based on site level incidence (n = 87). Statistical significance of P < 0.05 is marked in bold with a *.

| A. Virus incidence and prevalence |  |  |  |  | B. Pearson Correlations |  |  |  |  |  |  |
| --- | --- | --- | --- | --- | --- | --- | --- | --- | --- | --- | --- |
| Virus | Sample incidence | % sample incidence | site incidence | % site incidence | Virus | IAPV | DWV | SBV | ABPV | BQCV | CBPV |
| IAPV | 6 | 2% | 6 | 7% | IAPV | 1.00 |  |  |  |  |  |
| DWV | 92 | 36% | 47 | 55% | DWV | -0.03 | 1.00 |  |  |  |  |
| SBV | 65 | 25% | 36 | 42% | SBV | 0.14 | 0.02 | 1.00 |  |  |  |
| ABPV | 1 | >1% | 1 | 1% | ABPV | -0.03 | -0.12 | -0.09 | 1.00 |  |  |
| BQCV | 97 | 38% | 52 | 60% | BQCV | **0.22\*** | 0.17 | 0.16 | 0.09 | 1.00 |  |
| CBPV | 20 | 8% | 12 | 14% | CBPV | -0.11 | 0.03 | -0.07 | -0.04 | -0.02 | 1.00 |

Abreviations: Acute bee paralysis virus (ABPV), Black queen cell virus (BQCV), Chronic bee paralysis virus (CBPV), Deformed wing virus (DWV), Israeli acute paralysis virus (IAPV), and Sacbrood virus (SBV).

An average of 1.09 viruses were detected in each sample (SE = 0.058), thus the probability of fitting a Poisson distribution was 0.009. An average of 1.79 viruses were detected at each site (SE = 0.11), thus the probability of fitting a Poisson distribution was 0.006. This provides evidence against independent infection by the viruses assessed at both levels (D’Alvise et al., 2019). While DWV and SBV incidence was positively associated with BQCV, none of these were significantly correlated at the 95% confidence level. The only positive significant pairwise correlation was between IAPV and BQCV (*P* = 0.04).

### Effects of apiary management and landscape context on virus prevalence

We observed a positive relationship between regional bee density and BQCV as well as SBV; regional bee density was the only variable included in the best-fit models for these two viruses (Table 2). In contrast, we found that mite levels and julian date were the terms included in the best-fit model for DWV. For each virus, the full model included a positive influence of mites and regional density on disease prevalence, and a more variable, much less predictive negative influence of hive movement and a positive influence of no management on disease incidence. Each of the most prevalent viruses was found in both commercial and hobbyist apiaries, and in agricultural, mixed-use residential, and urban landscapes. Bee virus incidence differed by apiary management style for DWV (χ^2^ = 28.90, df = 2, *P* < 0.001), SBV (χ^2^ = 11.45, df = 2, *P* = 0.003), BQCV (χ^2^ = 4.65, df = 2, *P* = 0.10), CBPV (χ^2^ = 6.01, df = 2, *P* = 0.049) (Fig. 3). There was significantly higher incidence of DWV at sites without apiaries, many of which were located in urban community gardens, and a few in semi-natural roadside environments. There was higher incidence of SBV and BQCV at commercial apiaries (Fig. 3)

**Table 2.**
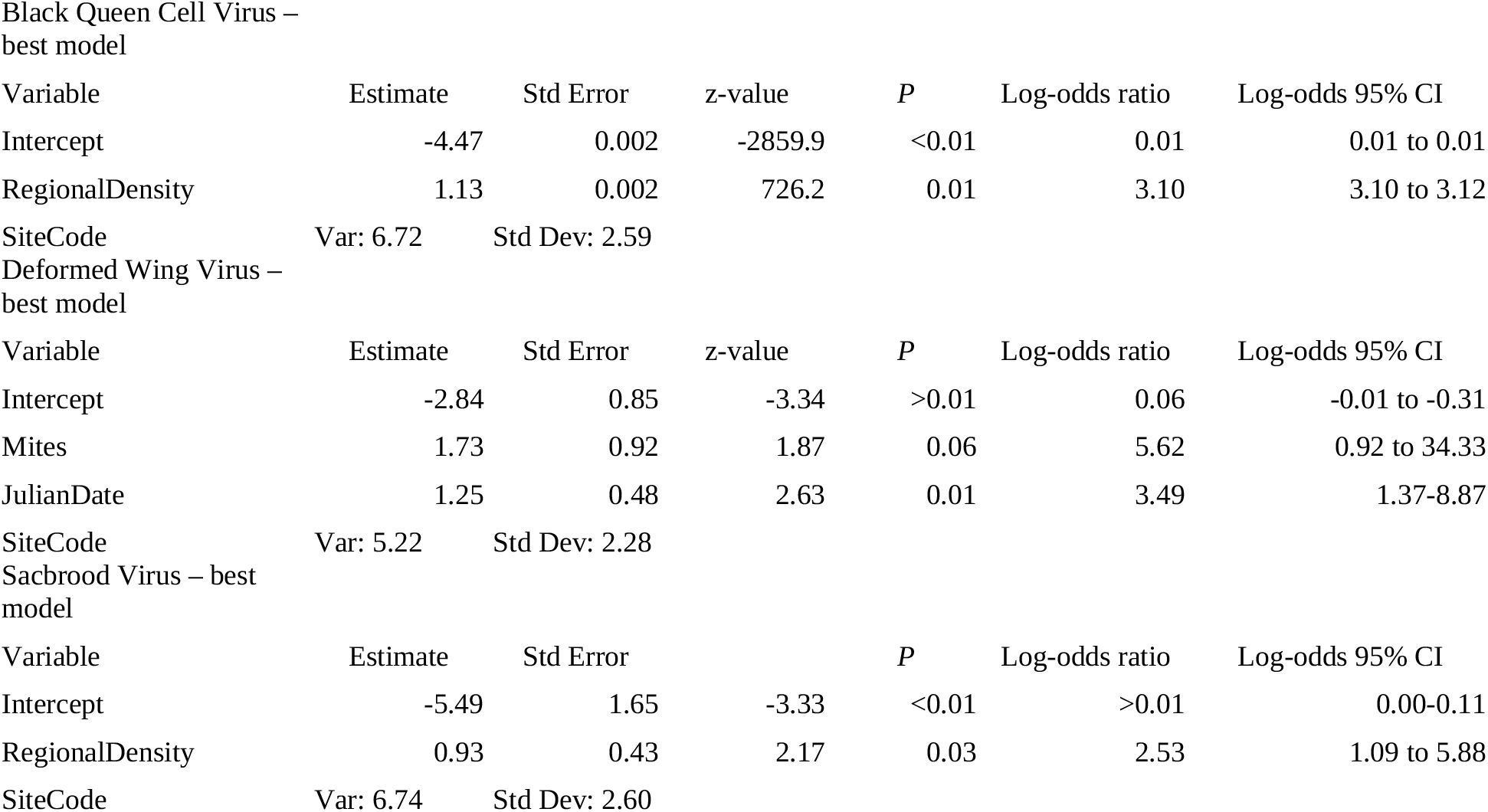
Best logistic regression mixed models for BQCV, DWV, SBV incidance. Top models were selected by AIC.

**Figure 2.**
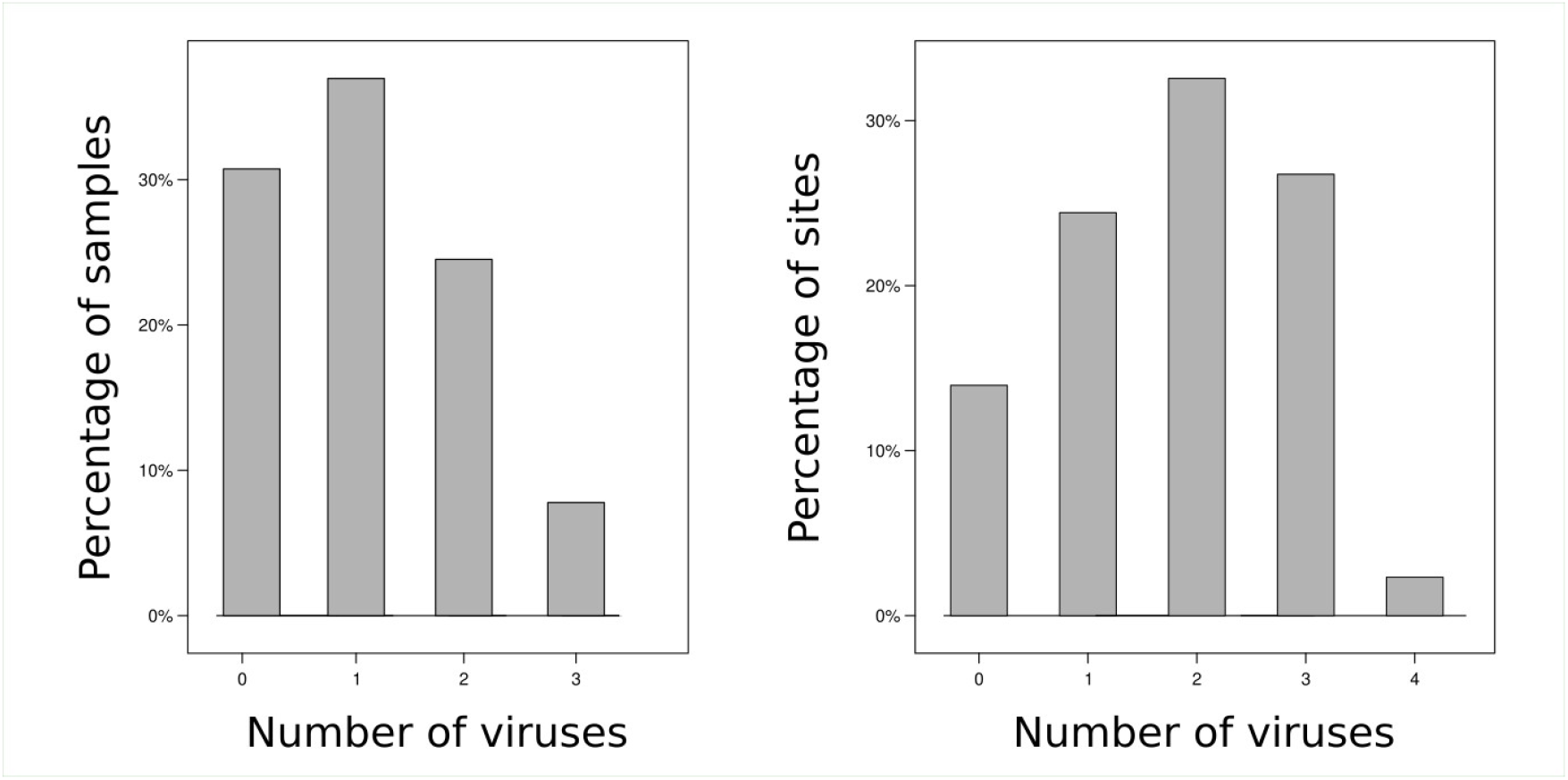
Number of viruses detected in (A) samples (n = 257) and (B) sites (n = 86)

**Figure 3.**
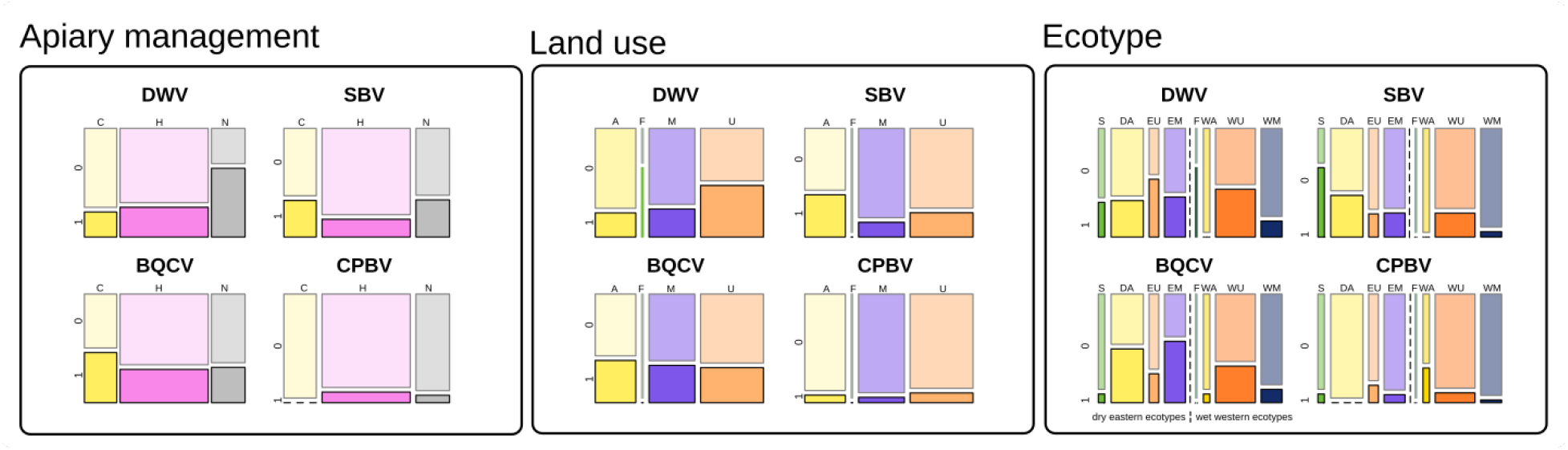
Mosaic plots show the number of positive (1) versus negative (0) tests for each virus across A. apiary management, i.e. commercial (n=54 tests), hobbyist (n=147), and non-apiary locations (n=56 tests) and B. land use, i.e. agriculture (n=69), forested (n=3 tests), mixed-use residential (n=78 tests), and urban (n=107 tests) and ecosystem type, (i.e. steppe, dryland agricultural, east-side urban, east-side mixed residential, cascades forest, west-side agricultural, west-side urban, and west-side mixed residential.

DWV and SBV incidence varied based on surrounding land use (χ^2^ = 17.47, df = 3, *P* = 0.001) and (χ^2^ = 15.06, df = 3, *P* = 0.002), respectively, while BQCV and CPBV did not (Fig. 4). DWV incidence was higher in urban and forested locations, compared to agricultural and mixed-use residential areas. SBV was highest in agricultural locations, followed by urban areas, and lowest in forested and mixed use residential areas (Figs. 1, 4).

#### Spatial autocorrelation among viruses

We assessed the role of additional spatial autocorrelation in our virus dataset using spatial regression, and did not find evidence of local spatial processes significantly influencing the distribution of the viruses. We applied the Moran’s test on regression residuals in preparation to fit a spatially lagged regression model, but did not observe sufficient spatial autocorrelation to proceed. The DWV moran’s I statistic standard deviate was 1.31 (*P* = 0.19), BQCV standard deviate was 1.42 (*P* = 0.15), and SBV standard deviate was 0.03 (*P* = 0.98).

## Discussion

Our study shows that regional apiary density and mites increased the incidence of common bee viruses, and disease-specific aspects of virus transmission ecology determined the best predictors to explain the prevalence of the three common viruses. DWV was observed more frequently in urban landscapes, and best predicted by mite levels, while SBV and BQCV were best predicted by regional bee density. While SBV was observed more frequently in agricultural landscapes and commercial apiaries, BQCV was common in cities with high bee density and in agricultural landscapes. DWV can be transmitted by mites, and mite treatment practices are somewhat more variable amongst hobbyists than commercial apiaries (Chen and Siede, 2007; Grozinger and Flenniken, 2019; Yang and Cox-Foster, 2007). SBV is not often associated with mites, but rather nurse bees spread the virus as they tend and remove infected larvae (Chen and Siede, 2007). SBV transmission is especially likely during the warm season, when commercial pollination of crops is underway, and while colonies are rearing susceptible brood. The high density of bees in large apiaries increases the chances of transmitting pathogens (Goulson and Hughes, 2015). Additional virus specific factors relating to virus transmissibility, such as reproduction number, may also mediate spread. For example, a less transmissible virus with a lower reproduction number may require a higher density of hosts to spread through a region.

BQCV, DWV, and SBV incidence exhibited similar patterns as other studies generally, although local sampling of commercial apiaries in high density bee regions have exhibited higher rates of virus incidence. Several studies of virus occurrence in commercial agricultural regions of Argentina, Germany, Turkey, and the United States (BQCV and DWV) have observed 90-100% incidence of common viruses (Alger et al., 2019; Cagirgan et al., 2020; D’Alvise et al., 2019; Murray et al., 2019). However, each of the three sporadically observed viruses from this study were also only observed occasionally in other North American studies, but in some other world regions, these three viruses are much more common. A Turkish study recently observed ABPV in 13 out of 15 colonies sampled, for example (Cagirgan et al., 2020).

We also observed evidence of synergistic effects between viruses, or shared influence of disease risk factors, leading to non-independent infection rates between viruses at the sample and colony level. While this pattern was observed overall, based on a higher than expected mean number of viruses per colony, significant correlation between viruses was only observed for IAPV and BQCV; correlations between SBV, DWV, and BQCV were not significant at the colony level. This analysis was used to investigate virus co-occurence between individual bees, and while distributions did not depart from Poisson distribution overall, spearman correlations in virus intensity were observed, indicating potential synergistic effects (D’Alvise et al., 2019).

Mites can transmit DWV, IAPV, and other pathogens to honey bees, and mite treatment can slow the spread of viruses. For example, experimental application of acaricide treatments in an experimental study was followed by a decrease in DWV titer as mites were brought under control (Locke, 2012). Our study did not observe an influence of mite treatment on the incidence of any of viruses, however; most apiaries use chemical treatment to control mites, however, so there was little variability in this factor. Yet, mite presence observed by beekeepers in the survey was the strongest predictor of DWV incidence, supporting the idea that mite treatment is a powerful tool to combat DWV spread in honey bees. Disease treatment styles varied more between hobbyist than commercial beekeepers, and study participants may be less variable than hobbyist beekeepers at large given their participation in beekeepers associations.

Virus incidence differed based on surrounding land use. When we split various land use categories by ecosystem type, based largely on the East-West precipitation gradient combined with surrounding land use in our study extent, the common viruses seemed much more common in eastern dryland agriculture and eastern mixed-use residential compared to western agriculture and mixed-use residential. Mixed-use residential was comprised by more exurban agriculture or rangeland on the eastern side of the Cascades Mountains, and more coniferous forest on the western side of the Cascades Mountains. Precipitation may have some direct influence on environmental contamination and transmission rates, but factors associated with commercial pollination and agriculture likely also contribute to the perceived differences.

While differences in virus incidence between land use types were observed, past studies suggest these patterns may not be consistent. For example, samples of 26 honey bee hives from near Lincoln, Nebraska, USA found no difference in the prevalence of DWV, BQCV, IAPV, and SBV between urban and agricultural landscapes (Olgun et al., 2020). Landscapes included in our surveys included regions with flowering crops (e.g. canola, apples, pears, and vegetable seed crops) that rely heavily on pollination from mobile apiaries. The contrast between extensive, commercially pollinated agricultural land use, cities with strong apiary communities, and coniferous forest rich natural and suburban landscapes likely generated the patterns we observed.

Our study shows mite monitoring and treatment may be help combat virus transmission between honey bees, especially in landscapes with a high density of apiaries. The spread and intensification of bee viruses is thought to be a major factor in increasing honey bee losses, and more attention and awareness of infectious diseases in apiculture could reduce virus spread. As colony losses remain high, but beekeeping continues to increase in popularity, understanding regional patterns of disease incidence and the mechanisms that underlie them are critical.

## Acknowledgments

We thank C. Delgado, M. Combs, G. Gregory, and O. Hopipa for assisting with virus testing. This project was funded by Western SARE grant (SW18-031).

## Citations

Alger, S.A., Burnham, P.A., Boncristiani, H.F., Brody, A.K., 2019. RNA virus spillover from managed honeybees (Apis mellifera) to wild bumblebees (Bombus spp.). PLoS ONE 14, e0217822. https://doi.org/10.1371/journal.pone.0217822

Aubert, M.F.A., European Commission, Directorate General for Research, Directorate E--Biotechnologies, A., Food, 2011. Virology and the honey bee. Northern Bee Books, Mytholmroyd, UK.

Bailey, L., 1969. The multiplication and spread of sacbrood virus of bees. Annals of Applied Biology 63, 483–491.

Bailey, L., Ball, B.V., Perry, J., 1981. The prevalence of viruses of honey bees in Britain. Annals of Applied Biology 97, 109–118.

Bailey, L., Gibbs, A., Woods, R., 1963. Two viruses from adult honey bees (Apis mellifera Linnaeus). Virology 21, 390–395.

Bailey, L., Perry, J., 1982. The diminished incidence of Acarapis woodi (Rennie)(Acari: Tarsonemidae) in honey bees, Apis mellifera L.(Hymenoptera: Apidae), in Britain. Bulletin of Entomological Research 72, 655–662.

Bailey, L., Woods, R., 1977. Two more small RNA viruses from honey bees and further observations on sacbrood and acute bee-paralysis viruses. Journal of General Virology 37, 175–182.

Boncristiani, H., Li, J., Evans, J.D., Pettis, J., Chen, Y., 2011. Scientific note on PCR inhibitors in the compound eyes of honey bees, Apis mellifera. Apidologie 42, 457–460.

Brutscher, L.M., McMenamin, A.J., Flenniken, M.L., 2016. The buzz about honey bee viruses. PLoS Pathogens 12, e1005757.

Burnham, A.J., McLaughlin, F., Burnham, P.A., Lehman, H.K., 2019. Local honey bees (Apis mellifera) have lower pathogen loads and higher productivity compared to non-local transplanted bees in North America. Journal of Apicultural Research 58, 694–701. https://doi.org/10.1080/00218839.2019.1632150

Cagirgan, A.A., Yildirim, Y., Usta, A., 2020. Phylogenetic analysis of deformed wing virus, black queen cell virus and acute bee paralysis viruses in Turkish honeybee colonies. Medycyna Weterynaryjna 76, 6437–2020. https://doi.org/10.21521/mw.6437

Chen, Y.P., Siede, R., 2007. Honey Bee Viruses, in: Advances in Virus Research. Elsevier, pp. 33–80. https://doi.org/10.1016/S0065-3527(07)70002-7

Cornman, R.S., Tarpy, D.R., Chen, Y., Jeffreys, L., Lopez, D., Pettis, J.S., vanEngelsdorp, D., Evans, J.D., 2012. Pathogen Webs in Collapsing Honey Bee Colonies. PLoS ONE 7, e43562. https://doi.org/10.1371/journal.pone.0043562

Cox-Foster, D.L., Conlan, S., Holmes, E.C., Palacios, G., Evans, J.D., Moran, N.A., Quan, P.-L., Briese, T., Hornig, M., Geiser, D.M., 2007. A metagenomic survey of microbes in honey bee colony collapse disorder. Science 318, 283–287.

D’Alvise, P., Seeburger, V., Gihring, K., Kieboom, M., Hasselmann, M., 2019. Seasonal dynamics and co-occurrence patterns of honey bee pathogens revealed by high-throughput RT-qPCR analysis. Ecol Evol 9, 10241–10252. https://doi.org/10.1002/ece3.5544

Di Prisco, G., Cavaliere, V., Annoscia, D., Varricchio, P., Caprio, E., Nazzi, F., Gargiulo, G., Pennacchio, F., 2013. Neonicotinoid clothianidin adversely affects insect immunity and promotes replication of a viral pathogen in honey bees. Proceedings of the National Academy of Sciences 110, 18466–18471.

Di Prisco, G., Pennacchio, F., Caprio, E., Boncristiani Jr, H.F., Evans, J.D., Chen, Y., 2011. Varroa destructor is an effective vector of Israeli acute paralysis virus in the honeybee, Apis mellifera. Journal of General Virology 92, 151–155.

Dolezal, A.G., Carrillo-Tripp, J., Judd, T.M., Allen Miller, W., Bonning, B.C., Toth, A.L., 2019. Interacting stressors matter: diet quality and virus infection in honeybee health. R. Soc. open sci. 6, 181803. https://doi.org/10.1098/rsos.181803

Doublet, V., Labarussias, M., de Miranda, J.R., Moritz, R.F.A., Paxton, R.J., 2015. Bees under stress: sublethal doses of a neonicotinoid pesticide and pathogens interact to elevate honey bee mortality across the life cycle: Pesticide-pathogen interactions in honey bees. Environ Microbiol 17, 969–983. https://doi.org/10.1111/1462-2920.12426

Dynes, T.L., Berry, J.A., Delaplane, K.S., Brosi, B.J., 2019. Reduced density and visually complex apiaries reduce parasite load and promote honey production and overwintering survival in honey bees 16.

Goulson, D., Hughes, W.O.H., 2015. Mitigating the anthropogenic spread of bee parasites to protect wild pollinators. Biological Conservation 191, 10–19. https://doi.org/10.1016/j.biocon.2015.06.023

Grozinger, C.M., Flenniken, M.L., 2019. Bee Viruses: Ecology, Pathogenicity, and Impacts. Annu. Rev. Entomol. 64, 205–226. https://doi.org/10.1146/annurev-ento-011118-111942

Locke, B., 2012. Host-Parasite Adaptations and Interactions Between Honey Bees, Varroa Mites and Viruses 76.

Loope, K.J., Baty, J.W., Lester, P.J., Wilson Rankin, E.E., 2019. Pathogen shifts in a honeybee predator following the arrival of the Varroa mite. Proc. R. Soc. B. 286, 20182499. https://doi.org/10.1098/rspb.2018.2499

Maori, E., Lavi, S., Mozes-Koch, R., Gantman, Y., Peretz, Y., Edelbaum, O., Tanne, E., Sela, I., 2007. Isolation and characterization of Israeli acute paralysis virus, a dicistrovirus affecting honeybees in Israel: evidence for diversity due to intra-and inter-species recombination. Journal of General Virology 88, 3428–3438.

Murray, E.A., Burand, J., Trikoz, N., Schnabel, J., Grab, H., Danforth, B.N., 2019. Viral transmission in honey bees and native bees, supported by a global black queen cell virus phylogeny. Environ Microbiol 21, 972–983. https://doi.org/10.1111/1462-2920.14501

Olgun, T., Everhart, S.E., Anderson, T., Wu-Smart, J., 2020. Comparative analysis of viruses in four bee species collected from agricultural, urban, and natural landscapes. PLOS ONE 21.

Tentcheva, D., Gauthier, L., Bagny, L., Fievet, J., Dainat, B., Cousserans, F., Colin, M.E., Bergoin, M., 2006. Comparative analysis of deformed wing virus (DWV) RNA in Apis mellifera and Varroa destructor. Apidologie 37, 41–50.

Tentcheva, D., Gauthier, L., Zappulla, N., Dainat, B., Cousserans, F., Colin, M.E., Bergoin, M., 2004. Prevalence and seasonal variations of six bee viruses in Apis mellifera L. and Varroa destructor mite populations in France. Applied and environmental microbiology 70, 7185–7191.

White, G.F., United States., Bureau of Entomology., United States., Department of Agriculture., 1913. Sacbrood, a disease of bees. U.S. Dept. of Agriculture, Bureau of Entomology, Washington, D.C.

Yang, X., Cox-Foster, D., 2007. Effects of parasitization by Varroa destructor on survivorship and physiological traits of Apis mellifera in correlation with viral incidence and microbial challenge. Parasitology 134, 405.

